# Application of aptamers improves CRISPR-based live imaging of plant telomeres

**DOI:** 10.1101/2020.05.05.078246

**Authors:** Solmaz Khosravi, Patrick Schindele, Evgeny Gladilin, Frank Dunemann, Twan Rutten, Holger Puchta, Andreas Houben

## Abstract

Development of live imaging techniques for providing information how chromatin is organized in living cells is pivotal to decipher the regulation of biological processes. Here, we demonstrate the improvement of a live imaging technique based on CRISPR/Cas9. In this approach, the sgRNA scaffold is fused to RNA aptamers including MS2 and PP7. When the dead Cas9 (dCas9) is co-expressed with chimeric sgRNA, the aptamer-binding proteins fused to fluorescent protein (MCP-FP and PCP-FP) are recruited to the targeted sequence. Compared to previous work with dCas9:GFP, we show that the quality of telomere labelling was improved in transiently transformed *Nicotiana benthamiana* using aptamer-based CRISPR-imaging constructs. Labelling is influenced by the copy number of aptamers and less by the promoter types. The same constructs were not applicable for labelling of repeats in stably transformed plants and roots. The constant interaction of the RNP complex with its target DNA might interfere with cellular processes.

**Highlight:** Aptamer-based CRISPR imaging: an opportunity for improving live-cell imaging in plants

## Introduction

The 3D organization of the genome is involved in the regulation of various genomic functions including gene expression, transcription, DNA replication, and repair (Misteli, 2007). Different strategies have been developed to monitor the dynamics of defined genomic loci in living cells (Chen *et al.*, 2013; Fujimoto *et al.*, 2016; Lindhout *et al.*, 2007; Robinett *et al.*, 1996; Saad H. *et al.*, 2014). Most recently, the clustered regularly interspaced short palindromic repeats (CRISPR)/CRISPR associated protein 9 (Cas9) based strategy has extensively been used mostly in non-plant species for live imaging. The first applications of CRISPR/Cas for live-cell imaging in plant (Dreissig *et al*., 2017; Fujimoto and Matsunaga, 2017) and non-plant cells (Chen *et al.*, 2013) was based on fluorescent proteins directly fused to deactivated Cas9 (dCas9). Different dCas9 orthologues from *Streptococcus pyogenes* and *Staphylococcus aureus* successfully label telomeres in transiently transformed *Nicotiana benthamiana* leaves (Dreissig *et al.*, 2017; Fujimoto and Matsunaga, 2017). Accordingly, it was shown that the locations of telomeres are in the periphery of the nucleus and dynamic positional changes of telomeres up to ± 2 µm were reported (Dreissig *et al.*, 2017).

Indirect labelling of dCas9 with the SunTag method resulted in 19 fold brighter signals in mammalian cell cultures in comparison to GFP-fused dCas9 (Tanenbaum *et al.*, 2014). However, this method like directly labelled dCas9 does not have the possibility of multi-targeting of genomic regions. For this purpose, different variants of dCas9 which have specific cognate gRNA were combined to label different genomic regions (Dreissig *et al.*, 2017; Esvelt *et al.*, 2013; Ma *et al.*, 2015). To improve the efficiency of imaging and also the capacity of dCas9 for multi-targeting of different regions at the same time, other methods for indirect labelling of dCas9 were adapted including BIFC (Hong *et al.*, 2018; Tanenbaum *et al.*, 2014), Aio-Casilio (Zhang and Song, 2017) and RNA-aptamer-based methods (Fu *et al.*, 2016; Ma *et al.*, 2016; Qin *et al.*, 2017; Shao *et al.*, 2016; Wang *et al.*, 2016). CRISPR-based live-cell imaging methods are reviewed in (Khosravi *et al.*, 2020; Wu *et al.*, 2019).

Among the improved indirect labelling methods, aptamer-based methods are used in mammalian cell cultures to target telomeric and other genomic regions. Aptamers are short RNA oligos which can be detected by specific RNA binding proteins (Urbanek *et al.*, 2014). Aptamer-based imaging methods are based on three components including dCas9, sgRNA in which the aptamer sequence is integrated and the aptamer binding protein which is fused to the fluorescent protein (Fu *et al.*, 2016; Ma *et al.*, 2016; Shao *et al.*, 2016). In plants, aptamers have been used for CRISPR/Cas9 targeted gene regulation with effector proteins like transcription activation domains, acetyltransferase or methyltransferase which were fused to the aptamer binding protein (Lee *et al.*, 2019; Selma *et al.*, 2019). The copy number of aptamers determines the number of effector proteins enriched in the targeted region. However, no application of CRISPR live-cell imaging based on aptamers is reported in plants yet.

In this research, we developed a CRISPR life imaging method based on the application of MS2 and PP7 aptamers for targeting telomers in transiently transformed *N. benthamiana*. We investigate whether the copy number of aptamers, sgRNA scaffold changes and promoter type affect labelling efficiency of target sequences. However, the same method was not successful for constant labelling of chromosome regions in stably transformed plants (*N. benthamiana* and *A. thaliana*) and roots (*Daucus carota*), suggesting that a continuous interaction of the RNP complex with target sequences might interfere with the progression of the cell cycle and plant development.

## Materials and Methods

### Plasmid construction

#### Expression of dCas9 driven by different promoters

To establish a three-component aptamer-based labelling method, dCas9 under the control of a ubiquitin parsley promoter was indirectly labelled with aptamer binding proteins (MS2 or PP7) fused to fluorescent proteins. The 35S promoter was amplified with *Eco*RI-35S-f1 and r1 primers flanking with an *Eco*RI recognition site from pCCNCEN (Supplementary Table 1). Then it was digested with *Eco*RI and cloned to linearized pDe-Sp-dCas9 GentR with *Eco*RI, which had another *Eco*RI site in the backbone removed before by site-directed mutation. The same method was used for substitution of the ubiquitin parsley promoter with a RPS5A promoter. The isolation of RPS5A was done by PRS5A-FWD and REV primers from the pGPTV-BAR (Supplementary Table 1). The XVE inducible promoter was generated with primers (Cas9-XVE-F, XVE-Lexa-A-R, XVE-Lexa-A-F and LexA-Cas9-R (Supplementary Table 1) containing homologous flanks for further Gibson Assembly into the pDe-Sp-dCas9 GentR. The pER8-v3 plasmid was used for generation of the XVE inducible promoter (Zuo *et al.*, 2000) (Supplementary Table 1). According to (Dreissig *et al.*, 2017), a pChimera expression gRNA vector in combination with a dCas9-eGFP expression vector was used as a control vector to target telomeres.

#### Insertion of aptamer sequences into the sgRNA scaffold

For aptamer-mediated imaging, sgRNA expression vectors were created either harbouring one MS2 aptamer sequence each in the tetraloop and stem-loop 2 of the *S. pyogenes* sgRNA backbone (Konermann et al. 2015) or three PP7 aptamer sequences only in the tetraloop of the *S. pyogenes* sgRNA backbone additionally comprising an A-U pair flip and stem extension (Shechner et al. 2015). In case of MS2, the vector pDS2.0-MS2 was synthesized comprising the respective sgRNA under control of the AtU6-26 promoter together with the codon-optimized MS2 binding protein cds joined to a 3’ SV40 NLS by a 3x GGGGS linker under control of the ZmUbi-1 promoter. In case of PP7, the respective sgRNA and codon-optimized PP7 binding protein cds also harboring a 3’ SV40 NLS were synthesized and subcloned via restriction digestion and ligation into pDS2.0-MS2 creating pDS2.0-PP7. *Bsm*BI restriction sites downstream of the aptamer binding protein cds were used for in-frame cloning of a 3-fold fusion of either eGFP or mRuby2. For this purpose, the respective cds were amplified from pSIM24-eGFP and pcDNA3-mRuby2 (www.addgene.com) with primers (MS2(NLS)-GFP#1-f, GFP#1-linker1-r, linker1-GFP#2-f, GFP#2-linker2-r, linker2-GFP#3-f, GFP#3-nos_ter-r or MS2(NLS)-mRuby#1-f, mRuby#1-linker1-r, linker1-mRuby#2-f, mRuby#2-linker2-r, linker2-mRuby#3-f, mRuby#3-nos_ter-r) adding homologous flanks for subsequent Gibson Assembly into the linearized pDS2.0-MS2 or pDS2.0-PP7 similar as previously described (Dreissig et al. 2017) creating pDS2.0-MS2/PP7-3xeGFP/3xmRuby2 (Supplementary Table 1).

#### Changing the sgRNA scaffold

An MS2 aptamer-harboring sgRNA additionally comprising an A-U flip and stem extension (Chen *et al.*, 2013) was synthesized and subcloned into pDS2.0-MS2-eGFP/mRuby2. For this purpose, pDS2.0-MS2-eGFP/mRuby2 was amplified with primers (pDS2.0-ΔsgRNA-r, pDS2.0-ΔsgRNA-f) deleting the sgRNA and the synthesized sgRNA was amplified with primers (sgRNA2.0-MS2-flip/ext-f, sgRNA2.0-MS2-flip/ext-r) adding overhangs for subsequent Gibson Assembly into the linearized backbone (Supplementary Table 1).

#### Altering the copy number of aptamers

To change the copy number of aptamers, pDS.2.0-MS2+3xeGFP gRNA expression vector was used. To delete one of MS2 copy numbers, pDS.2.0-MS2+3xeGFP was double digested with *Age*l and *Msc*I restriction enzymes and then was ligated to annealed primers Apta2-FWD and Apta2-Rev flanked by *Age*l overhang (Supplementary Table 1). Annealing of primers was done by mixing 2 μl of each primer (100 pM) in the total volume of 50 μl double distilled water and incubation at 95 °C. Colony PCR was performed by SS42 and Apta2-Rev2 primers under following conditions: 95°C for 5 min, 30x (95°C for 30 sec, 58°C for 30 sec, 72°C for 30 sec.), 72°C 5 min. Positive clones were confirmed by sequencing with the SS42 primer (Supplementary Table 1). To increase the copy number of aptamer sequences, a pDS2.0-MS2-eGFP/mRuby2 sgRNA expression vector was used. First, according to Qin et al., 2017 a sgRNA scaffold harbouring 16 MS2 aptamers was synthesized and subcloned into pDS2.0-MS2-eGFP/mRuby2. For this purpose, pDS2.0-MS2-eGFP/mRuby2 was digested with *Bsm*BI and *Age*I for sgRNA deletion and the synthesized sgRNA was digested with *Bsa*I and *Age*I for subsequent ligation into the linearized pDS2.0-MS2-eGFP/mRuby2 creating pDS2.0-16xMS2-eGFP/mRuby2.

#### Designing protospacers for targeting different genomic regions

The protospacer design was performed with the help of DeskGen (https://www.deskgen.com/). Each protospacer sequence was selected based on the PAM sequence of SpCas9 and synthesized as primer oligos with appropriate overhangs at 5’ ends for cloning into the pDS2.0-MS2:3xeGFP/mRuby (Supplementary Table 1). Then, the pDS2.0-MS2:3xeGFP/mRuby was subcloned to dCas9 expression vector by Gateway cloning. The dCas9 expression vector carries a gentamycin resistant marker for selection of stably transformed plants. The telomere protospacer was designed based on *Arabidopsis*-type telomere repeat sequence 5′-(TTTAGGG)(n)-3′. *Arabidopsis*-type centromere-specific protospacers were designed based on centromeric satellite consensus sequences (Supplementary Table 1).

#### Plant material and transformation

All imaging constructs were separately transformed to *Agrobacterium tumefaciens* GV3101. For carrot transformation, *A. rhizogenes* 15843 was used. Agrobacteria were cultured overnight in LB medium containing spectinomycin (100 mg/l^-1^) and rifampicin (50 mg/l^-1^) for transient transformation of *N. benthamiana* according to (Phan and Conrad, 2016). Additionally, a *N. benthamiana* line expressing CFP-histone H2B was used (Martin *et al.*, 2009). For the telomeric repeat binding protein 1 fused to GFP (TRB1-GFP), Agrobacteria were cultured in LB medium containing kanamycin (100 mg/l^-1^) and rifampicin (50 mg/l^-1^) (Schrumpfová *et al.*, 2014). For co-transformation experiments, bacterial cultures with the same OD600 were mixed in a 1:1 ratio. Stable transformation of *N. benthamiana, Daucus carota* (cultivars Blanche, Yellowstone and Rotin) and *A. thaliana* (var. Colombia) with dCas9:2xMS2:GFP constructs were performed via leaf samples, A. rhizogenes-based hairy root transformation and floral dip method according to (Clemente, 2006), (Dunemann *et al.*, 2019) and (Martin *et al.*, 2009), respectively. PCR and real-time PCR were performed for putative transgenic plants using primers specific for dCas9 and GFP to confirm the presence and expression of T-DNA fragments (Supplementary Table 1).

#### Immunostaining and fluorescence *in situ* hybridization (FISH)

Sampling for immunostaining was performed three days after infiltration of *N. benthamiana*. Briefly, a piece of leaf tissue with the size of ∼1 cm^2^ was excised and chopped in 0.5 ml chromosome isolation buffer (Doležel *et al.*, 2007) and then filtrated through a 35 μm nylon mesh with subsequent centrifugation onto microscopic slides with a CytoSpin3 (Shandon) at 400 rpm for 5 min. To confirm the specificity of signals CRISPR imaging and FISH were combined. The intensity of CRISPR signals was increased by in direct immunostaining using a 1:2,500 diluted Dylight 488-labelled GFP mouse monoclonal antibody (cat. 200-341-215, Rockland) according to (Ishii *et al.*, 2015). Detection of *Arabidopsis*-type telomeres via FISH was performed with a 5’Cy5-labelled probe (5’GGGTTTAGGGTTTAGGGTTT). Immuno-FISH was performed as described by (Ishii *et al.*, 2015). Immunostaining against dCas9, was performed with a DyLight 550-labelled SpCas9 mouse monoclonal antibody (cat. NBP2-52398R, Novus Biological).

#### Proteasome inhibitor test

The plants were kept on MS medium containing 50, 100 or 150 µM MG-132 (Serva) under dark condition at room temperature for 16 h.

#### Microscopy

Micrographs were captured using an epifluorescence microscope (Olympus BX61) equipped with a cooled charge coupled device (CCD) camera (Orca ER; Hamamatsu). Images were collected from at least 10 nuclei per experiment and then analyzed with ImageJ. For live-cell imaging, a confocal laser scanning microscope (LSM780, Carl Zeiss) was used. To detect fluorescence signals *in vivo*, a piece of infiltrated leaf was cut and with the use of 40x NA 1.2 water objective nuclei with clear signals were tracked for 20 minutes. 488-nm laser line was used for excision of GFP and emission was detected over a range of 490-540 nm.

#### Analysis of telomere signals

To measure the labelling efficiency of telomeres, 20 nuclei were imaged for each construct by epifluorescent microscope. The number of telomere signals per nucleus was determined and the mean value was calculated. To evaluate the signal/background noise, the maximum signal intensity was divided by minimum signal intensity rising from the background using the ImageJ software. The mean value was calculated from three measurements in each nucleus.

To study the movement of telomeres, telomere tracking was performed for 5 nuclei and was based on time-laps z stacks from IMARIS 8.0 (Bitplane). The adjustments to calculate the coordinates (*x, y, z*) of each telomere and also measuring the inter-telomere distances was based on Dreissig et al. (2017). To assess true displacements of telomeres over time, global movements of nuclei have to be computationally eliminated. For this purpose, 3D point clouds of telomere mass centres for all subsequent time steps (*t>0*) were rigidly registered to the reference system of coordinates defined by the first time step (*t=0*) using absolute orientation quaternions (Horn, 1987). To quantify the intranuclear telomere motion, the mean square distance (*MSD*) of telomeres relatively to their initial position (t=0) was calculated as

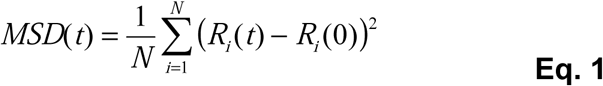

where *Ri(t)* is the radius vector of the *i-th* registered telomere in the reference system of coordinates at the time point *t>0*.

## Results

### Optimizing live imaging of telomeres with aptamer-based CRISPR/dCas9 imaging vectors

The application of fluorescent proteins directly fused to dCas9 resulted in the labelling of ∼27 telomeres of 72 expected signals in 2C nuclei of *N. benthamiana* (Dreissig et al. 2017). To improve the labelling efficiency, we established RNA aptamer-based CRISPR/dCas9 imaging constructs for plants. The three-component constructs (called dCas9:2xMS2:GFP and dCas9:3xPP7:GFP) encode dCas9 of *S. pyogenes*, an *Arabidopsis* telomere-specific sgRNA with integrated aptamer sequences (2x MS2 or 3x PP7) and aptamer coat proteins fused to three copies of fluorescent proteins (tdMCP:GFP or tdPCP:GFP binding to MS2 or PP7 aptamers, respectively) (Fig. 1A, B). In addition, a dCas9:2xMS2 construct with a 3x mRuby-tagged coat protein (called dCas9:2xMS2:mRuby) was prepared (Supplementary Fig. S1).

**Figure 1.**
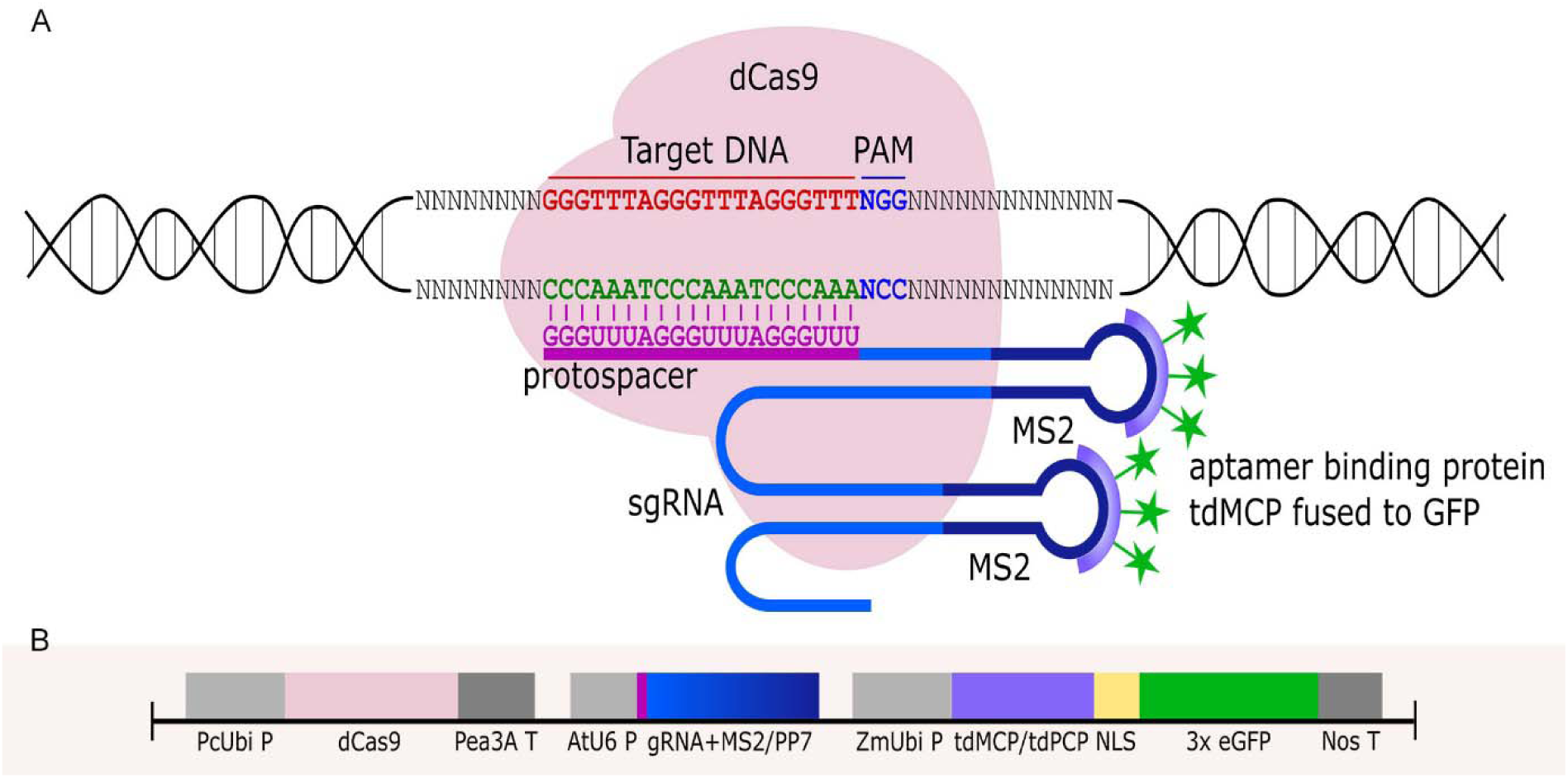
RNA aptamer-based CRISPR/dCas9 imaging of telomere repeats. (a) Schemata depicting the components of the aptamer-based CRISPR labelling method: 1) dCas9 from *S. pyogenes*, 2) MS2 or PP7 aptamers (here only MS2 is shown) which are integrated into the sgRNA scaffold. 3) RNA binding protein (tdMCP or tdPCP) fused to fluorescent protein (3x eGFP) which recognizes aptamers. Protospacer designed to target *Arabidopsis*-type telomere DNA sequence. (b) Structure of the aptamer-based CRISPR imaging construct. dCas9 is driven by a ubiquitin promoter from parsley (PcUbi P), chimeric gRNA including aptamers (MS2/ PP7) are driven by the AtU6 promoter (AtU6 P), aptamer binding proteins fused to a fluorescent protein (tdMCP/ tdPCP) with the help of nuclear localization signal (NLS) are driven by a ubiquitin promoter from maize (ZmUbi P). Pea3A T and Nos T were used as terminators.

To compare the labelling efficiency of the newly designed constructs, *N. benthamiana* leaves were separately infiltered with both types of *Arabidopsis*-type telomere-specific dCas9-aptamer constructs (dCas9:2xMS2:GFP and dCas9:3xPP7:GFP) and the previously employed dCas9:GFP reporter (Dreissig *et al.*, 2017). Both types of aptamer-based constructs successfully labelled telomeres in interphase nuclei (Fig. 2A, B). In average, 48 and 37 signals were recognized by dCas9-2xMS2:GFP and dCas9-3xPP7:GFP, respectively (Fig. 2D). In contrast, the application of dCas9:GFP resulted in ∼28 CRISPR-based signals which is consistent with earlier research (Dreissig *et al.*, 2017) (Fig. 2C, D). The lower number of detected signals than the expected could be due to clustering of some telomeres or not all telomeres were detectable by the applied imaging constructs. Notably, the accumulation of GFP signals in the nucleolus, which was always observed by application of dCas9:GFP was not found in nuclei labelled with both types of dCas9-aptamer constructs (Fig. 2A, B, C).

**Figure 2.**
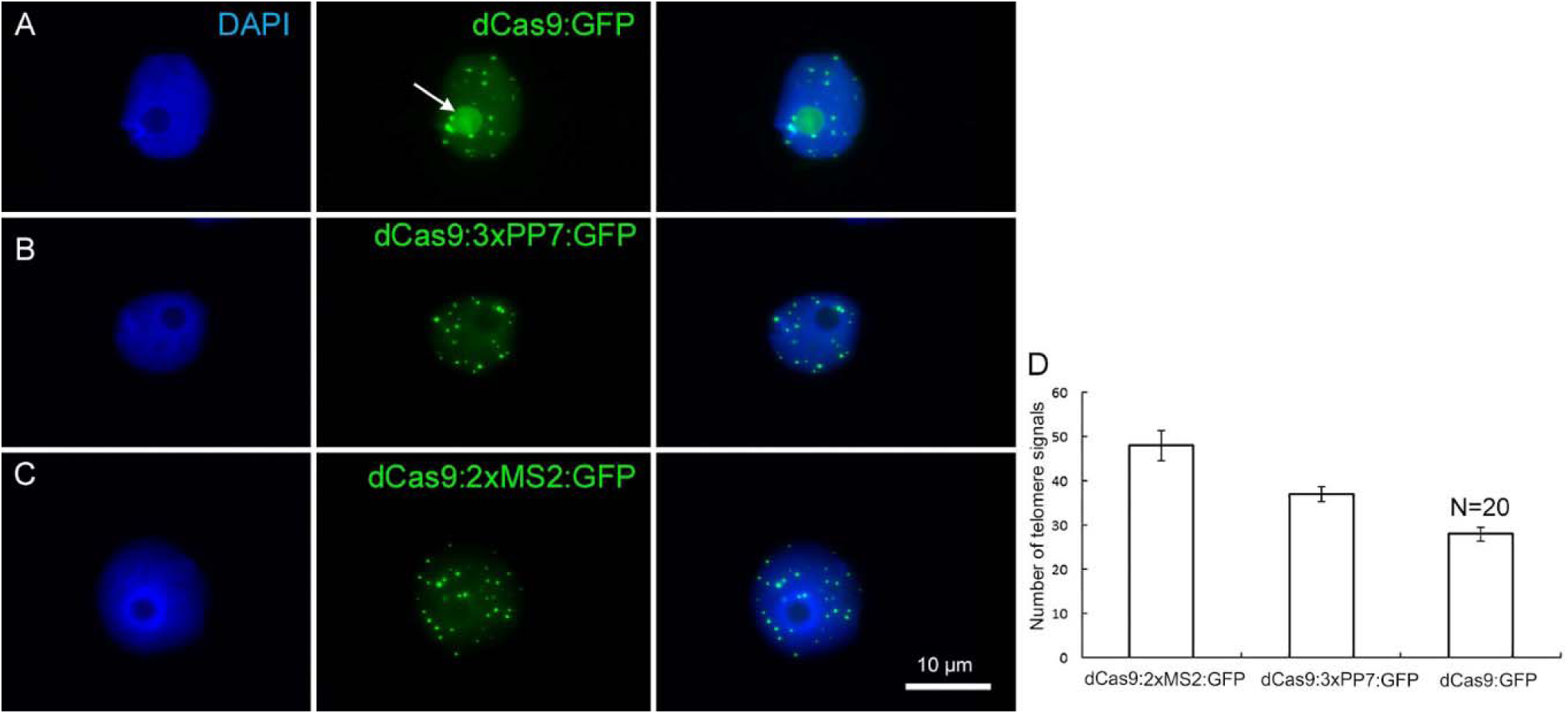
Live imaging of telomeres in *N. benthamiana* leaf cells during interphase by CRISPR/dCas9. The distribution of telomeres recognized by (a) dCas9:GFP (b) dCas9:3xPP7:GFP and (c) dCas9:2xMS2:GFP. Note, aptamer-based imaging constructs (dCas9:3xPP7:GFP and dCas9:2xMS2:GFP) did not label nucleoli, while the application of dCas9:GFP does (nucleolus shown with white arrow). Nuclei are counterstained with DAPI. (d) Diagram showing the efficiency of indirectly and directly labelled dCas9 for targeting telomeric regions. The number of telomere signals was determined based on 20 nuclei per construct. dCas9 indirectly labelled either with MS2 or PP7 aptamers shows more telomeres (p<0.05).

As a negative control, the transformation of *N. benthamiana* with partial constructs carrying dCas9:GFP without target-specific gRNA or pMS2:mRuby targeting telomers without the dCas9 component was performed. For both, a nonspecific labelling of nuclei was found (Fig. 3A, B). After co-transformation with both partial constructs, overlapping telomere-like signals of green and red fluorescence were found due to the presence of all components required for CRISPR imaging of telomeres (Fig. 3C).

**Figure 3.**
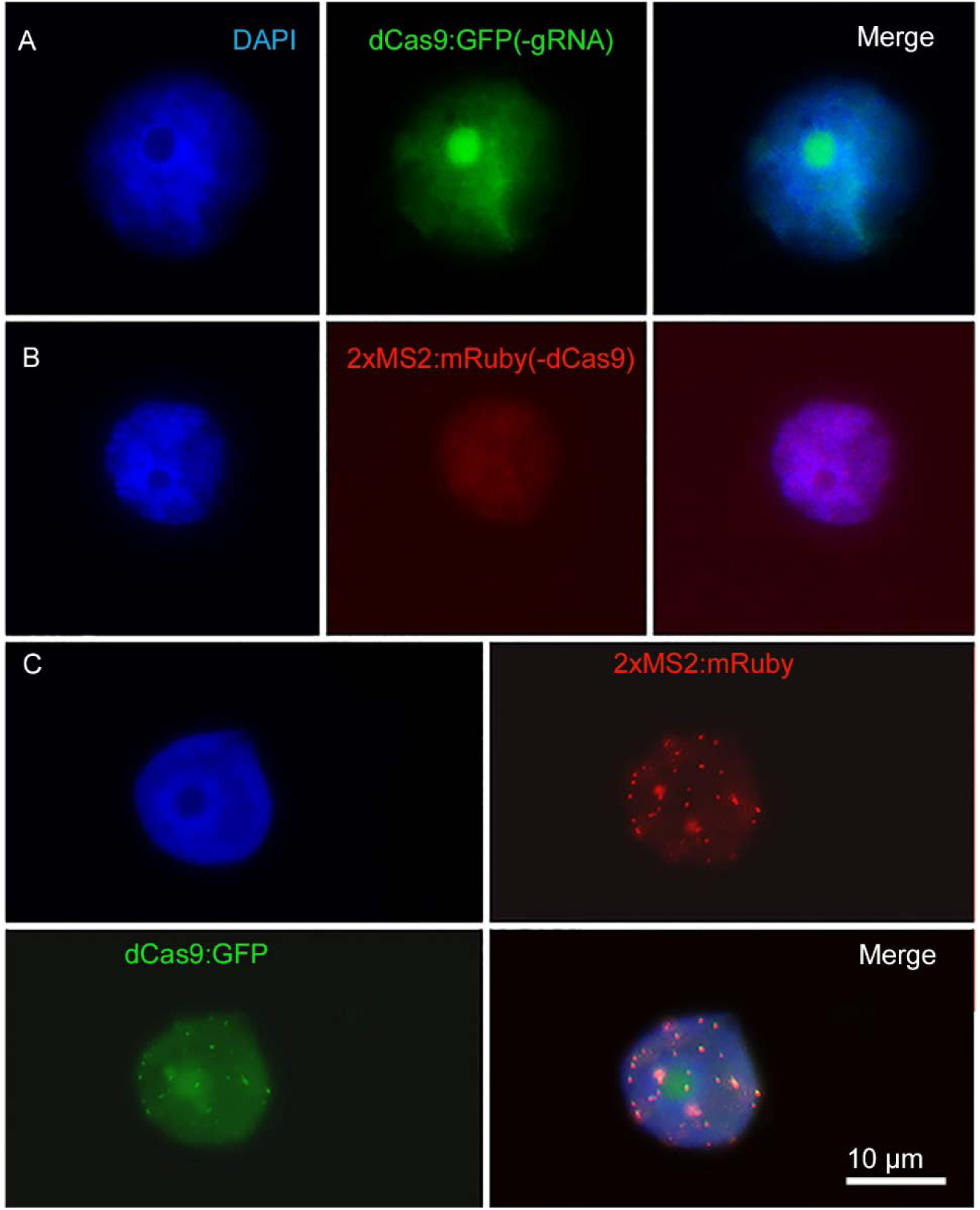
Negative control with partial constructs carrying (a) dCas9:GFP without gRNA or (b) 2xMS2:3xmRuby targeting telomers without dCas9. (c) Co-transformation of *N. benthamiana* leaves with both partial dCas9:GFP and 2xMS2:3xmRuby constructs resulted in labelling of telomeres, while no telomere-like signals were found after transformation with either partial construct (a, b). Nuclei are counterstained with DAPI.

To confirm the target specificity of the observed telomere-like signals, FISH with a labelled telomere-specific probe was performed after CRISPR imaging. All dCas9:2xMS2:GFP signals co-localized with FISH signals, demonstrating the target specificity of the aptamer-based imaging approach (Fig. 4A). However, the labelling efficiency of CRISPR was less than FISH as only 78% and 75% of FISH signals colocalized with dCas9:2xMS2:GFP and dCas9:3xPP7:GFP signals, respectively (Fig. 4B). Co-expression of dCas9:2xMS2:mRuby with TRB1 and telomeric dCas9:2xMS2:GFP with CFP labelled histone H2B (Martin *et al.*, 2009) showed that the aptamer-based CRISPR imaging method can also be successfully combined with fluorescence-labelled proteins to study DNA-protein interactions (Supplementary Movie S1,S2).

**Figure 4.**
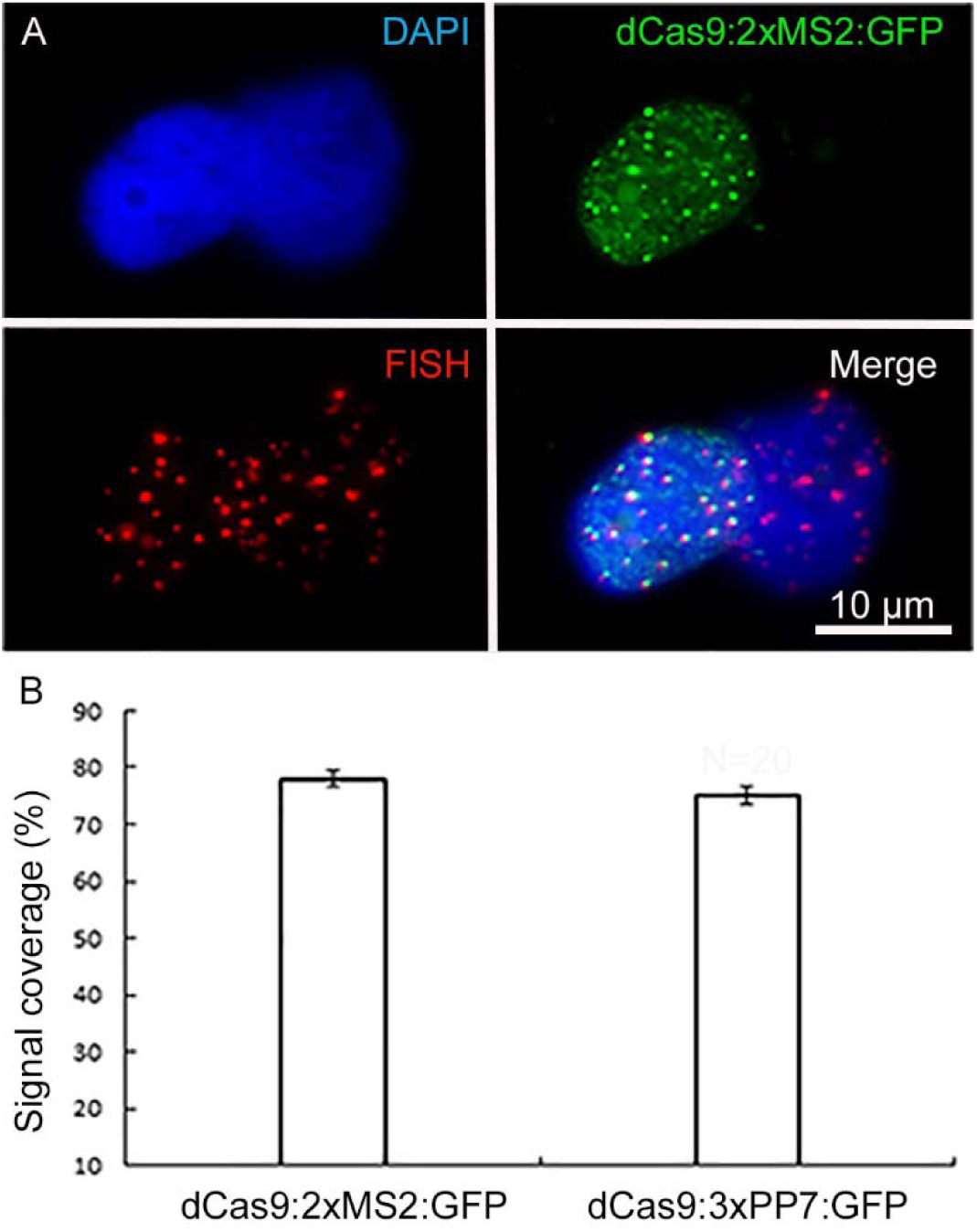
Confirming the target specificity of aptamer-based CRISPR imaging. (a) Immunofluorescence staining against dCas9:2xMS2:GFP combined with telomere-specific FISH. Nuclei are counterstained with DAPI (in blue). (b) Comparing the efficiency of both types of aptamer-based CRISPR imaging with FISH. Telomeric signals based on 20 isolated nuclei per each construct after ImmunoFISH. dCas9:2xMs2:GFP and dCas9:3xPP7:GFP recognized 78% and 75% of telomere signals identified by FISH, respectively (p<0.05).

To test whether the copy number of aptamers affects the labelling efficiency, we compared dCas9:MS2:GFP carrying 1, 2 or 16 copies of the MS2 aptamer. By reducing the aptamer copy number to 1, the number of observed signals reduced (Fig. 5A). 14 copies of MS2 did not result in enhanced telomere signals, instead strong background signals were produced (Fig. 5C).

**Figure 5.**
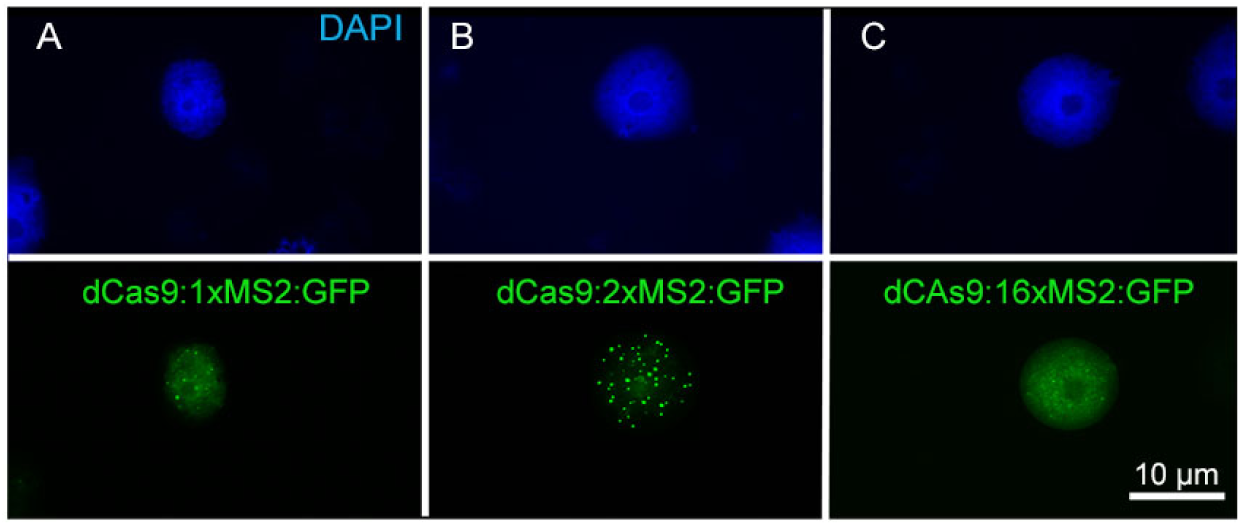
Effect of MS2 aptamer copy number of aptamer-based CRISPR imaging constructs on signal intensity. (a) dCas9:1xMS2, (b) dCas9:2xMS2 and (c) dCas9:16xMS2. The construct with two copies of MS2 revealed the best labelling of telomeres. Nuclei are counterstained with DAPI.

Because four sequential U nucleotides in the sgRNA stem-loop could be recognized as a transcription termination signal for the *A. thaliana* derived U6 pol-III promoter, a U to A substitution was performed and also the structure of sgRNA was changed by the insertion of an extension to improve the stability of sgRNA and its assembly with dCas9 according to (Supplementary Fig. S2). The U/A flip along with increasing the length of the sgRNA stem size did not result in a significant increase of telomere signal intensity and did not improve the signal/background noise ratio of telomere signals in *N. benthamiana* (Fig. 6A, B, C).

**Figure 6.**
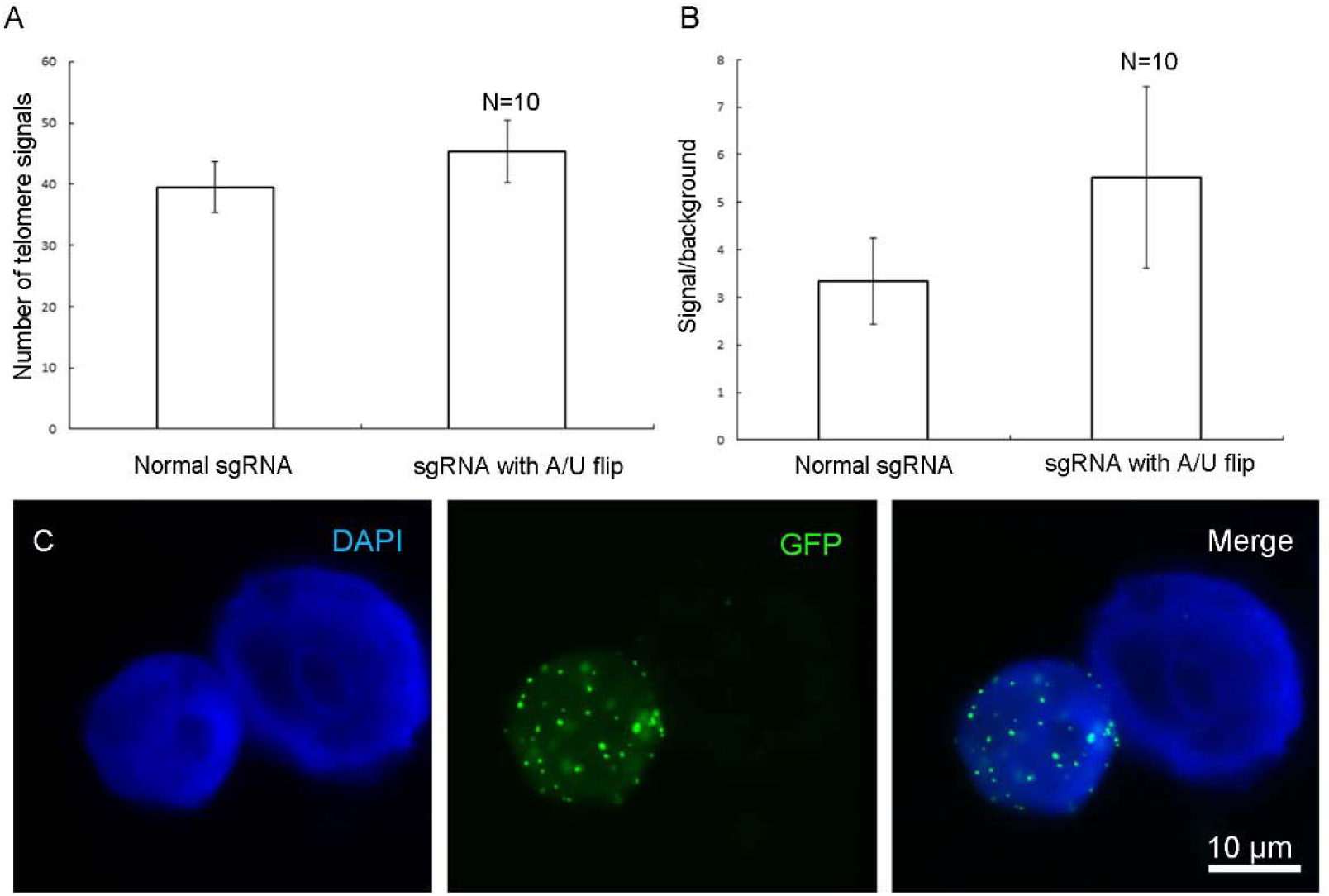
Effect of changing the sgRNA scaffold with a U/A flip and extension on quantity and quality of observed telomere signals. No significant change was observed in the terms of a) telomere number or b) signal/background noise ratio(p<0.05). c) Labelled telomeres by the vector which has the change in sgRNA scaffold. Measurements were performed based on data from 10 isolated nuclei.

### Comparing the effect of different promoters to express dCas9

Beside the ubiquitin promoter from parsley to drive the expression of dCas9 in *N. benthamiana*, we tested the Cauliflower mosaic virus (CaMV) 35S (Tepfer *et al.*, 2004), RPS5A (Weijers *et al.*, 2001) and the β-estradiol inducible promoter XVE (Zuo *et al.*, 2000). Changing the promoter in dCas9:2xMS2:GFP construct did not increase the number of observed telomere signals in comparison to the ubiquitin promoter (Fig. 7A). The 35S promoter led to a better signal/background noise ratio (Fig. 7B). After induction of the β-estradiol inducible XVE promoter, the same number of telomere signals was observed which was recognized by the construct driven by the ubiquitin promoter (Fig. 7A). The specificity of signals was approved by subsequent FISH with a telomere-specific probe (Supplementary Fig. S3A). Without induction, no telomere-specific signal was observed (Supplementary Fig. S3B).

**Figure 7.**
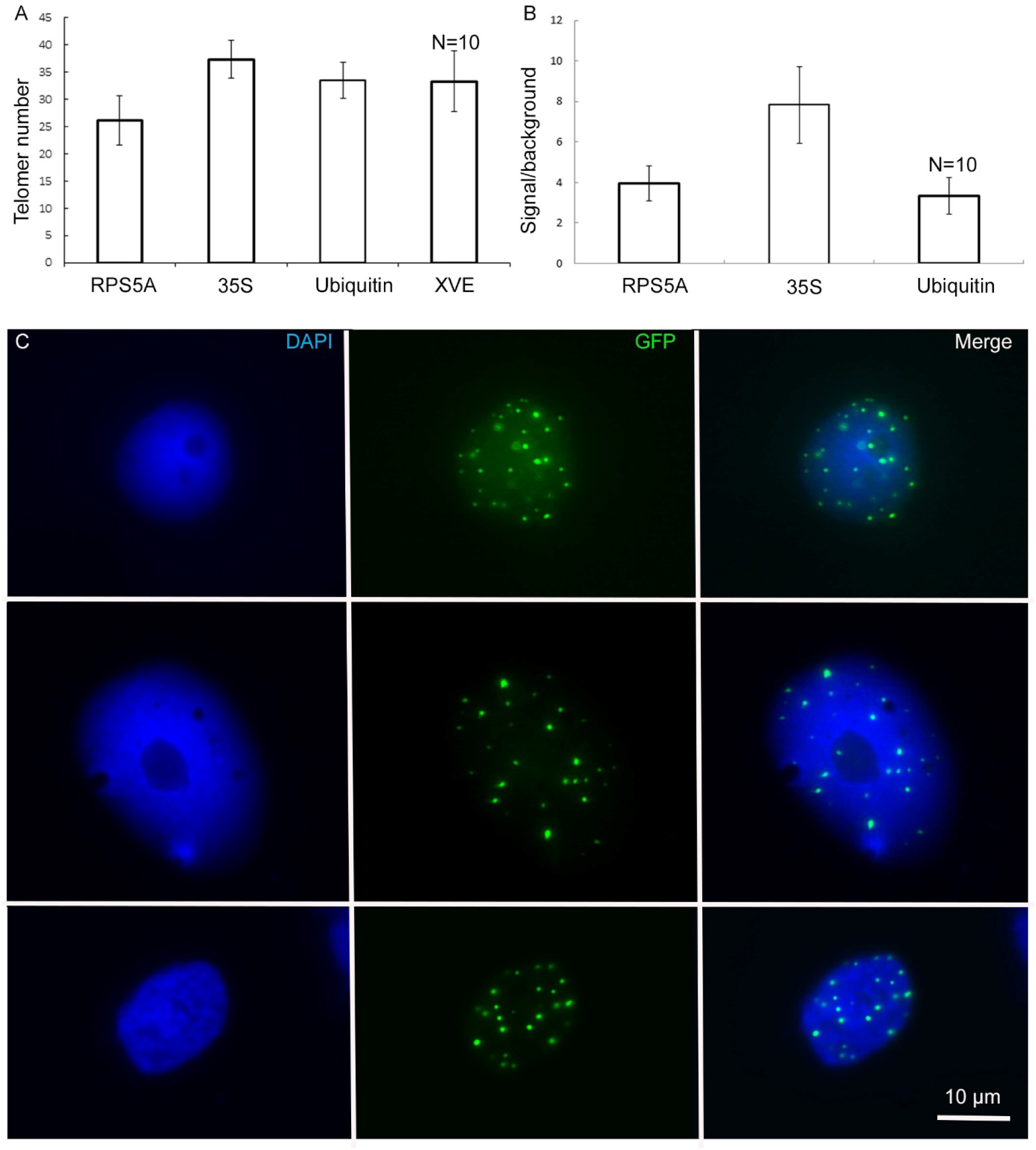
Effect of different promotors used for expression of dCas9 on the efficiency of telomere labelling. a) The expression of dCas9 by PRS5A promoter resulted in the recognition of a smaller number of telomeres compared to 35S and ubiquitin promoters. 35S promoter caused the better signal to background noise ratio. c) using XVE inducible promoter was as efficient as ubiquitin promoter regarding the number of labelled telomeres (p<0.05). Data obtained from 10 isolated nuclei per construct. Regardless of promoter type, dCas9 driven by d) RPS5A, e) 35S, f) XVE could label telomeric regions in *N. benthamiana*.

Comparison of dCas9 transcription driven by the XVE or ubiquitin promoter revealed that even weak dCas9 expression by XVE is sufficient to produce telomere-specific CRISPR-based signals (Supplementary Fig. S4). Regardless of the promoter type, telomeres showed similar dynamic and random movements (Fig. 8). To quantify these movements the mean square displacement (MSD) of telomeres was measured over a period of time. Calculating the changes of intratelomeric distance showed the minimum ±1 µm to maximum ±4 µm of changes for each type of promoter (Fig. 9). In summary, application of RNA-aptamers for CRISPR-based live-cell imaging increases the efficiency of telomere labelling in plant cells.

**Figure 8.**
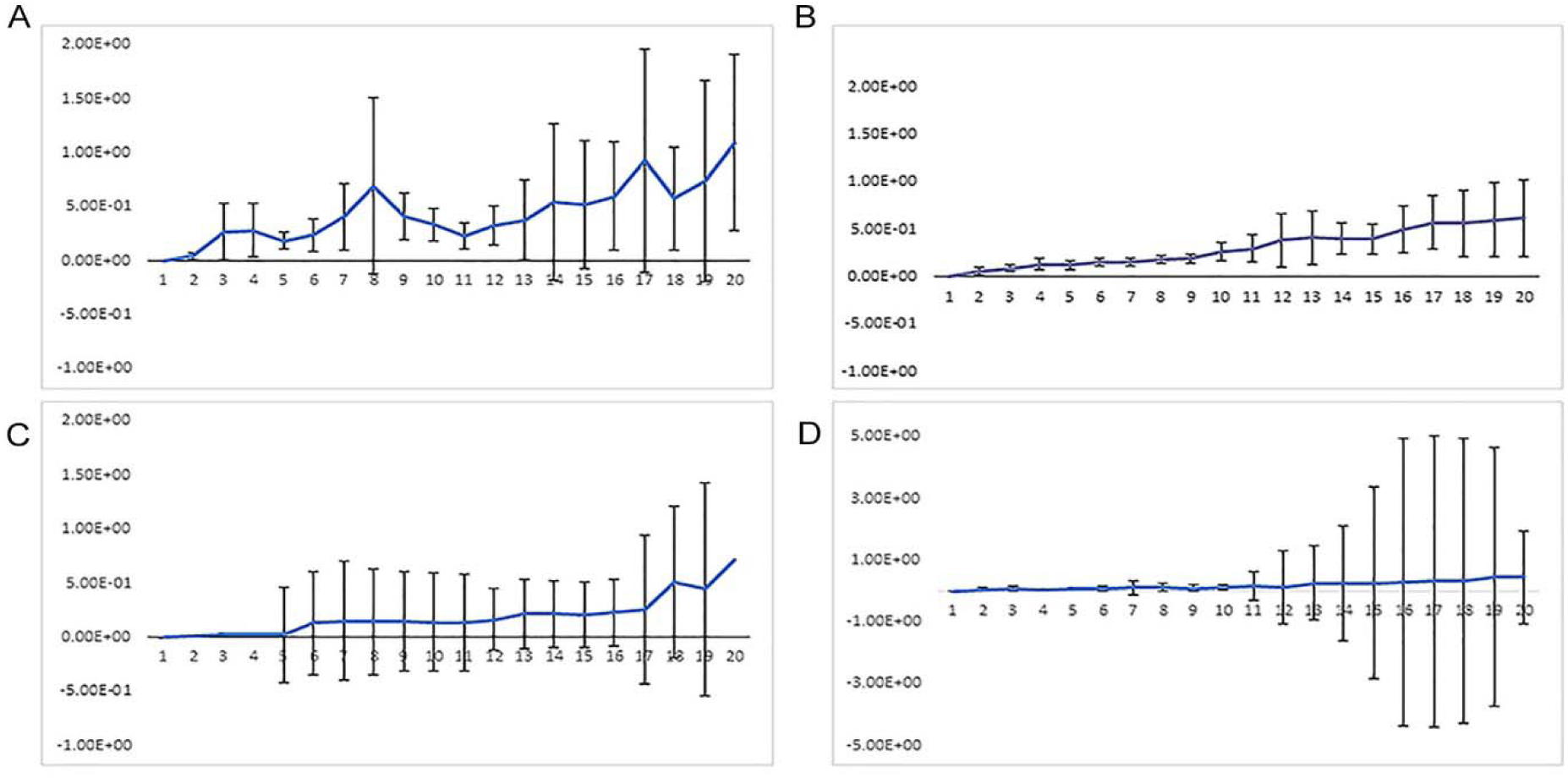
Comparing mean square distance (MSD in µm) of telomeres labelled by indirectly labelled aptamer-dCas9 which were under the control of a) 35S, b) RPS5a or c) ubiquitin promoters. d) Directly labelled dCas9, which was under the control of a ubiquitin promoter. Telomeres showed random movements regardless of promoter type and how dCas9 was labelled.

**Figure 9.**
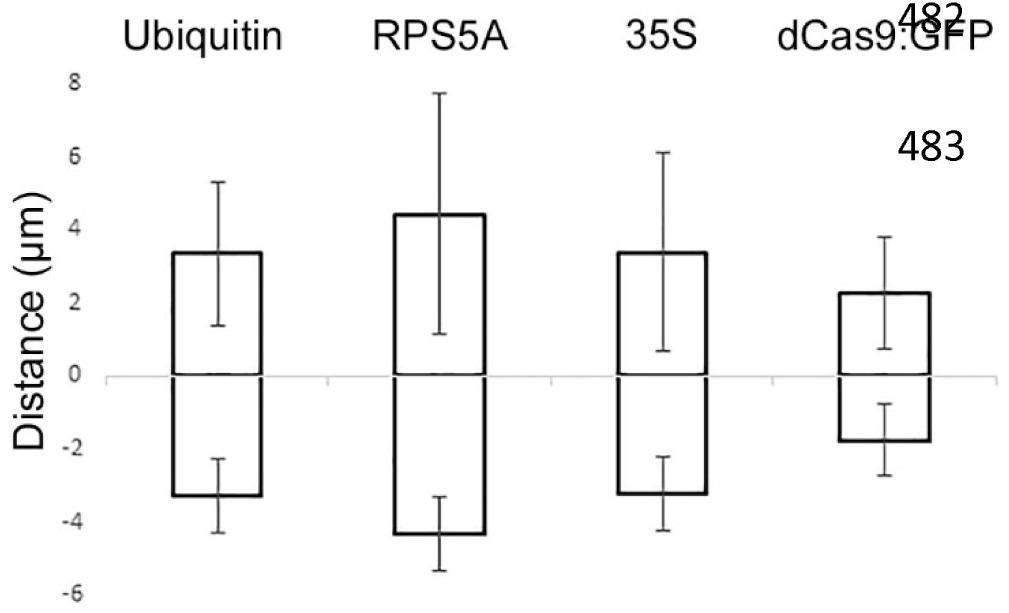
Measurement of inter-telomeric distance changes in nuclei transformed with three different indirectly labelled aptamer-dCas9 which were under the control of ubiquitin, RPS5a or 35S promoters and directly labelled dCas9 which was under the control of the ubiquitin promoter. Intra-telomeric distance changes vary between minimum ±1 µm to maximum ±4 µm.

### Application of CRISPR-imaging is limited in stably transformed plants

Stable transformation of *N. benthamiana, A. thaliana* plants and *D. carota* roots with the telomere-specific dCas9:2xMS2:GFP construct did not result in transgenic plants exhibiting GFP-labelled telomeres in living leaf or root cells, although the presence and expression of dCas9 and GFP genes were confirmed by PCR and real-time RT-PCR (data not shown, Supplement Table 2). Only transformation of *A. thaliana* with dCas9:2xMS2:GFP targeting centromeric regions resulted in few plants that showed some dot-like signals, however, the number and pattern of signals were atypical for interphase centromeres (Supplementary Fig. S5). In total, 141 selection marker resistant *A. thaliana* plants were screened for three different centromere imaging constructs by microscopy. Among them, 27 plants showed uniform labelling of nuclei and 9 plants showed dot-like signals. The dot-like signals were unstable and could not be detected in seedlings older than three weeks or subsequent generations (T3). Phenotype and seed setting of plants exhibiting dot-like signals were wild-type like. Among the three different protospacers used, only protospacer 1 and 2 produced signals. The same protospacer 1 was successfully used to label centromeres in fixed nuclei of *A. thaliana* with the help of CRISPR-FISH (Ishii *et al.*, 2019).

Plants that were transformed with dCas9:2xMS2:GFP under the control of an inducible promoter with a centromere- or telomere-specific protospacer revealed no target sequence-specific signals after induction with β-estradiol (Supplementary Table S2).

To test whether the disappearance of dot-like signals is caused by degradation of the dCas9 protein, transgenic plants were treated with different concentrations of the proteasome inhibitor MG-132. However, no dot-like signals were recovered. Additionally, the presence of dCas9 protein was confirmed by dCas9 immunostaining (Supplementary Fig. S6).

## Discussion

### Optimization of aptamer-based CRISPR imaging constructs

The application of MS2 and PP7 aptamers resulted in improved CRISPR imaging constructs instrumental to trace telomeres in transiently transformed *N. benthamiana*., Labelling efficiency, based on the mean value of signal numbers per nucleus, was increased up to 1.7 fold in comparison to dCas9:GFP. The number of individual telomere signals per nucleus was lower than expected though, which may be due to clustering of individual telomeres. Clustering of telomeres has been also observed in other organisms like *A. thaliana* (Fransz *et al.*, 2002), yeast and *Drosophila melanogaster* (Hozé *et al.*, 2013; Wesolowska *et al.*, 2013).

Despite the improved labelling of telomeres, the aptamer-based CRISPR imaging in *N. benthamiana* resulted in a labelling efficiency of 73 - 75% compared with FISH. In contrast, in human cell cultures, the number of telomeric signals obtained by CRISPR imaging was almost equal to the number of FISH signals (Chen *et al.*, 2013). The copy number difference of telomere repeats is unlikely the reason for this discrepancy because human telomeres are 5 to 15 kb (Moyzis *et al.*, 1988) while the telomeres in *N. benthami*ana are 60 to 160 kb long (Fajkus *et al*. 1995). Since a temperature of 37°C is required for optimal Cas9 activity (Xiang *et al.*, 2017), the temperature difference between plant (22°C) and mammalian cell cultures (37°C) might contribute to the observed labelling difference between mammalian and plant species.

While dCas9:GFP expressing cells showed background signals in nucleoli (Dreissig *et al.*, 2017), such background was absent from leaves expressing aptamer-containing reporter constructs. Nucleolar accumulation of dCas9 has been noted in other species like human cell cultures (Chen *et al.*, 2013). Likely, unspecific labelling of nucleoli was reduced because fluorescent proteins were not directly fused to dCas9.

Substitution of the ubiquitin promoter with the inducible XVE promoter caused a 5 fold decrease in expression of dCas9. However, changing the expression of dCas9 gene by application of XVE promoter did not result in a significant change in the number of observed telomere signals. In contrast, it is demonstrated that the low applied dosage of sgRNA in mammalian cell cultures affects the quality of CRISPR imaging signals (Chen *et al.*, 2013). The PRS5A promoter resulted in a lower number of telomere signals. This could be because PRS5A is more active in meristematic tissues rather than leaves, the tissue which was used for transient transformation (Winter *et al.*, 2007).

Increasing the number of MS2 aptamers to 16 copies did not enhance the efficiency of telomere labelling in *N. benthamiana*, although in human cell cultures increment of aptamer numbers up to 16 improved labelling (Qin *et al.*, 2017). Additionally, changing the sgRNA scaffold did not increase the quantity and quality of observed signals. In human cell cultures though, similar modifications increased the number of CRISPR-labelled telomeres and improved the signal/background noise (Chen *et al.*, 2013). Fujimoto and Matsunaga (2017) used sgRNA scaffold modifications (T to G change and A/U flip combined with UGCUG extension) within a CRISPR imaging construct to improve the signal to noise ratio of telomere labeling in transiently transformed *N. tabacum.* The different outcome reported here might be due to the different constructs used.

### Why does CRISPR imaging not work in stably transformed plants?

Our CRISPR imaging constructs which were successfully applied in transiently transformed *N. benthamiana* leaves could not be used to label defined sequences in stably transformed *N. benthamiana, A. thaliana* or *D. carota*. The same observation was made by (Fujimoto and Matsunaga, 2017) for GFP-fused dCas9 imaging constructs. Intriguingly, CRISPR-imaging of centromeric and telomeric repeats works fine on fixed nuclei and chromosomes of different plant and animal species (Deng *et al.*, 2015; Ishii *et al.*, 2019; Nemeckova *et al.*, 2019; Potlapalli *et al.*, 2020). The *in situ* imaging method CRISPR-FISH (also called REGEN-ISL) is based on a fluorescence-labelled two-part guide RNA with a recombinant Cas9 endonuclease complex. For both imaging methods, we used telomere- and centromere-specific gRNA and *A. thaliana* and *N. benthamiana*, subsequently (Ishii *et al.*, 2019), this work). Hence, our expectation was that the selected gRNA in combination with dCas9 should also work in stably transformed plants.

Why then did CRISPR imaging fail in stably transformed plants? In contrast to CRISPR-based editing, for CRISPR imaging a constant interaction of the RNP complex with the target DNA is a functional prerequisite. It is tempting to speculate that a permanent binding of the RNP complex with its target DNA interferes with processes required for plant development. The formation of R-loops, which is underlying the CRISPR/Cas mechanism, might hamper cellular processes. R-loops are three-stranded nucleic acid structures composed of a DNA-RNA hybrid and a displaced single-stranded DNA. R-loops have a role in transcription, chromatin modification, DNA damage response. Once the R-loop homeostasis is perturbed, it can lead to genome instability (Crossley *et al.*, 2019; Palmer, 2020). The R-loop distribution atlas of *A. thaliana* has shown that R-loop distribution patterns are relatively preserved during different developmental and environmental conditions (Xu *et al.*, 2020). Therefore, by imposing consistent formation of R-loops in targeted regions, CRISPR imaging constructs might change R-loop dynamics in defined genomic regions of stably transformed plants. Alternatively, the selected Cas9 variant of *S. pyogenes* is not suitable and further optimized Cas variants with higher efficiency could overcome this problem. A negative selection against CRISPR-imaging constructs in stably transformed plants at the transcript level is less likely because corresponding transcripts exist. In addition, uniform labelling of anti-Cas9 immunosignals was detected in transformed plants. Overcoming the discussed problem will also help to increase the efficiency of CRISPR-based editing in plants.

Taking advantage of the intrinsic stability of CRISPR guide RNA, (Wang *et al.*, 2019) used fluorescent ribonucleoproteins consisting of chemically synthesized fluorescent gRNAs and recombinant dCas9 protein for imaging in transfected living human lymphocytes. Live-cell fluorescent *in situ* hybridization (LiveFISH) allowed tracking of multiple chromosomal loci in lymphocytes. Whether the transient transformation of cells with fluorescent RNP complexes could become another option to label defined sequences in living plant cells remains to be demonstrated.

## Conclusions

A three-component labelling method using dCas9, PP7/MS2 aptamers and tdMCP:GFP/ tdPCP:GFP binding to MS2 /PP7 aptamers was successfully applied for labelling of telomers in transiently transformed *N. benthamiana*. The labelling efficiency of telomeres was increased and the background labelling noise in the nucleolus was reduced compared to previous work (Dreissig *et al.*, 2017). The copy number of aptamers used in the aptamer-based imaging construct is critical. The level of *dCas9* gene expression does not affect CRISPR imaging. The application of CRISPR/Cas9 for live-cell imaging in stably transformed plants, however, was not successful.

## Supplements

Movie supplement 1. Co-expression of dCas9:2xMS2:mRuby (red) and TRB1 (green) in *N. benthamina*. Co-localization of telomeric dCas9:2xMS2:mRuby and TRB1 shows that aptamer-based imaging construct can be also used for DNA-protein interaction studies. Movie supplement 2. Dynamic of telomeres targeted by indirectly labelled dCas9 with MS2 aptamer in *N. benthamiana* leaf nuclei expressing CFP-H2B.

Figure supplement 1. Different components of the aptamer-based labelling method: 1) dCas9 from *S. pyogenes*, 2) MS2 or PP7 aptamers (here only MS2 is shown) which are integrated into sgRNA scaffold. 3) RNA binding protein (tdMCP or tdPCP) fused to a fluorescent protein (mRuby) which recognizes aptamers.

Figure supplement 2. Changing the sgRNA scaffold with A/U flip (in red) and insertion of an extension (in green).

Figure supplement 3. Specificity control test by ImmunoFISH for the activity of the inducible XVE promoter. a) Isolated nuclei after treatment of leaves with β-estradiol show telomeric signals. Co-localization of dCas9:2xMS2:GFP and FISH signals show that the observed signals are telomeric specific. b) Nuclei isolated from β-estradiol-untreated leaves show a uniform labelling of nuclei.

Figure supplement 4. Real time expression of dCas9 expressed by ubiquitin and XVE promoters. dCas9 expression is much lower when it is driven by inducible XVE promoter compared to ubiquitin from parsley. Error bars are standard deviation.

Figure supplement 5. Selected nuclei of *A. thaliana* stably transformed with a centromere-specific dCas9:2xMS2:GFP construct exhibiting dot-like signals. a, b) Application of the centromere-specific protospacer 1 and 2, respectively. The number of signals was higher than expected.

Figure supplement 6. Immunostaining of dCas9 protein in isolated nuclei from leaf material of stably transformed *Arabidopsis* plants with dCas9:2xMS2:GFP targeting centromeric regions. a) Immunostaining of dCas9 in stably transformed Arabidopsis plants showed the dCas9 is not degraded. b) Immunostaining of isolated leaf nuclei from wild type *Arabidopsis* leaf nuclei did not result in signals which shows that the applied antibody against dCas9 is working specifically.

Table supplement 1. List of primers used in cloning steps and PCR reactions.

Table supplement 2. Summary of CRISPR live-cell imaging in stably transformed plants.

## Acknowledgements

We would like to thank Sabine Struckmeyer, Christine Helmold, Oda Weiß, Christin-Sophie Gäde and Sylvia Swetik for technical support, Steven Dreissig for scientific discussion and Christian Hertig for preparing schemata for the manuscript. We also thank Michael M. Goodin for providing *N. benthamiana* seeds expressing CFP-H2B, Bruno Müller for a plasmid containing the RPS5A promoter, Chua Nam Hai for the pER8 containing XVE promoter, Martina Dvořáčková and Jiri Fajkus for plasmid expressing TRB1-GFP. The work was funded by Deutsche Forschungsgemeinschaft (DFG) grant HO1779/28-1.

## References

Chen B, Gilbert LA, Cimini BA, Schnitzbauer J, Zhang W, Li GW, Park J, Blackburn EH, Weissman JS, Qi LS, Huang B. 2013. Dynamic imaging of genomic loci in living human cells by an optimized CRISPR/Cas system. Cell 155, 1479–1491.

Clemente T. 2006. *Nicotiana (Nicotiana tobaccum, Nicotiana benthamiana*). In: Wang K, ed. Agrobacterium Protocols, Vol. 1. Totowa, NJ: Hummana Press, 153–154.

Crossley MP, Bocek M, Cimprich KA. 2019. R-Loops as cellular regulators and genomic threats. Mol Cell 73, 398–411.

Deng WL, Shi XH, Tjian R, Lionnet T, Singer RH. 2015. CASFISH: CRISPR/Cas9-mediated in situ labeling of genomic loci in fixed cells. Proceedings of the National Academy of Sciences of the United States of America 112, 11870–11875.

Doležel J, Greilhuber J, Suda J. 2007. Estimation of nuclear DNA content in plants using flow cytometry. Nature Protocols 2, 2233–2244.

Dreissig S, Schiml S, Schindele P, Weiss O, Rutten T, Schubert V, Gladilin E, Mette MF, Puchta H, Houben A. 2017. Live-cell CRISPR imaging in plants reveals dynamic telomere movements. The Plant Journal 91, 565–573.

Dunemann F, Unkel K, Sprink T. 2019. Using CRISPR/Cas9 to produce haploid inducers of carrot through targeted mutations of centromeric histone H3 (CENH3). In: Grzebelus D, Barański R, eds. II International Symposium on Carrot and Other Apiaceae: ISHS Acta Horticulturae.

Esvelt KM, Mali P, Braff JL, Moosburner M, Yaung SJ, Church GM. 2013. Orthogonal Cas9 proteins for RNA-guided gene regulation and editing. Nature Methods 10, 1116.

Fajkus J, Kovařík A, mKrálovics R, Bezdek M. 1995. Organization of telomeric and subtelomeric chromatin in the higher plant *Nicotiana tabacum*. Molecular and General Genetics. 247, 633–638.

Fransz P, de Jong JH, Lysak M, Castiglione MR, Schubert I. 2002. Interphase chromosomes in Arabidopsis are organized as well defined chromocenters from which euchromatin loops emanate. Proceedings of the National Academy of Sciences of the United States of America 99, 14584–14589.

Fu Y, Rocha PP, Luo VM, Raviram R, Deng Y, Mazzoni EO, Skok JA. 2016. CRISPR-dCas9 and sgRNA scaffolds enable dual-colour live imaging of satellite sequences and repeat-enriched individual loci. Nat Commun 7, 11707.

Fujimoto S, Matsunaga S. 2017. Visualization of Chromatin Loci with Transiently Expressed CRISPR/Cas9 in Plants. Cytologia 82, 559–562.

Fujimoto S, Sugano SS, Kuwata K, Osakabe K, Matsunaga S. 2016. Visualization of specific repetitive genomic sequences with fluorescent TALEs in *Arabidopsis thaliana*. Journal of Experimental Botany 67, 6101–6110.

Hong Y, Lu G, Duan J, Liu W, Zhang Y. 2018. Comparison and optimization of CRISPR/dCas9/gRNA genome-labeling systems for live cell imaging. Genome Biology 19, 39.

Horn BKP. 1987. Closed-form solution of absolute orientation using unit quaternions. Journal of the Optical Society of America a-Optics Image Science and Vision 4, 629–642.

Hozé N, Ruault M, Amoruso C, Taddei A, Holcman D. 2013. Spatial telomere organization and clustering in yeast Saccharomyces cerevisiae nucleus is generated by a random dynamics of aggregation–dissociation. Molecular Biology of the Cell 24, 1791–1800.

Ishii T, Schubert V, Khosravi S, Dreissig S, Metje-Sprink J, Sprink T, Fuchs J, Meister A, Houben A. 2019. RNA-guided endonuclease - in situ labelling (RGEN-ISL): a fast CRISPR/Cas9-based method to label genomic sequences in various species. New Phytologist 222, 1652–1661.

Ishii T, Sunamura N, Matsumoto A, Eltayeb AE, Tsujimoto H. 2015. Preferential recruitment of the maternal centromere-specific histone H3 (CENH3) in oat (*Avena sativa* L.) × pearl millet (*Pennisetum glaucum* L.) hybrid embryos. Chromosome Research 23, 709–718.

Khosravi S, Ishii T, Dreissig S, Houben A. 2020. Application and prospects of CRISPR/Cas9-based methods to trace defined genomic sequences in living and fixed plant cells. Chromosome Research 28, 7–17.

Lee JE, Neumann M, Iglesias Duro D, Schmid M. 2019. CRISPR-based tools for targeted transcriptional and epigenetic regulation in plants. PLoS ONE 14, e0222778.

Lindhout BI, Fransz P, Tessadori F, Meckel T, Hooykaas PJJ, Zaal BJ. 2007. Live cell imaging of repetitive DNA sequences via GFP-tagged polydactyl zinc finger proteins. Nucleic Acids Res 35, e107.

Ma H, Naserib A, Reyes-Gutierreza P, Wolfec SA, Zhangb S, Pedersona T. 2015. Multicolor CRISPR labeling of chromosomal loci in human cells. PNAS 112, 3002–3007.

Ma H, Tu LC, Naseri A, Huisman M, Zhang S, Grunwald D, Pederson T. 2016. Multiplexed labeling of genomic loci with dCas9 and engineered sgRNAs using CRISPRainbow. Nat Biotechnol 34, 528–530.

Martin K, Kopperud K, Chakrabarty R, Banerjee R, Brooks R, Goodin MM. 2009. Transient expression in *Nicotiana benthamiana* fluorescent marker lines provides enhanced definition of protein localization, movement and interactions in planta. The Plant Journal 59, 150–162.

Misteli T. 2007. Beyond the Sequence: Cellular Organization of Genome Function. Cell 128, 787–800.

Moyzis RK, Buckingham JM, Cram LS, Dani M, Deaven LL, Jones MD, Meyne J, Ratliff RL, Wu JR. 1988. A highly conserved repetitive DNA sequence, (TTAGGG)n, present at the telomeres of human chromosomes. Proceedings of the National Academy of Sciences of the United States of America 85, 6622–6626.

Nemeckova A, Wasch C, Schubert V, Ishii T, Hribova E, Houben A. 2019. CRISPR/Cas9-based RGEN-ISL allows the simultaneous and specific visualization of proteins, DNA repeats, and sites of DNA replication. Cytogenetics Genome Research 159, 48–53.

Palmer L. 2020. The R-loop: An Additional Chromatin Feature for Gene Regulation in *Arabidopsis* Plant Cell 32, 785–786.

Phan HT, Conrad U. 2016. Plant-Based Vaccine Antigen Production. In: Brun A, ed. Vaccine Technologies for Veterinary Viral Diseases: Methods and Protocols. New York, NY: Springer New York, 35–47.

Potlapalli BP, Schubert V, Metje-Sprink J, Liehr T, Houben A. 2020. Application of Tris-HCl allows the specific labeling of regularly prepared chromosomes by CRISPR-FISH. Cytogenet Genome Research 160, 156–165.

Qin P, Parlak M, Kuscu C, Bandaria J, Mir M, Szlachta K, Singh R, Darzacq X, Yildiz A, Adli M. 2017. Live cell imaging of low- and non-repetitive chromosome loci using CRISPR-Cas9. Nature Communications 8, 14725.

Robinett CC, Straight AF, Li GW, Willhelm C, Sudlow G, Murray A, Belmont AS. 1996. In vivo localization of DNA sequences and visualization of large-scale chromatin organization using lac operator/repressor recognition. journal of cell biology 135, 1685–1700.

Saad H., Gallardo F., Dalvai M., Tanguy-le-Gac N., Lane D., K. B. 2014. DNA dynamics during early double-strand break processing revealed by non-intrusive imaging of living cells. PLoS Genet 10.

Schrumpfová PP, Vychodilová I, Dvořáčková M, Majerská J, Dokládal L, Schořová S, Fajkus J. 2014. Telomere repeat binding proteins are functional components of *Arabidopsis* telomeres and interact with telomerase. The Plant Journal 77, 770–781.

Selma S, Bernabé-Orts JM, Vazquez-Vilar M, Diego-Martin B, Ajenjo M, Garcia-Carpintero V, Granell A, Orzaez D. 2019. Strong gene activation in plants with genome-wide specificity using a new orthogonal CRISPR/Cas9-based programmable transcriptional activator. Plant Biotechnology Journal 17, 1703–1705.

Shao S, Zhang W, Hu H, Xue B, Qin J, Sun C, Sun Y, Wei W, Sun Y. 2016. Long-term dual-color tracking of genomic loci by modified sgRNAs of the CRISPR/Cas9 system. Nucleic Acids Research 44, e86.

Tanenbaum ME, Gilbert LA, Qi LS, Weissman JS, Vale RD. 2014. A protein tagging system for signal amplification in gene expression and fluorescence imaging. Cell 159, 635.

Tepfer M, Gaubert S, Leroux-Coyau M, Prince S, Houdebine L. 2004. Transient expression in mammalian cells of transgenes transcribed from the Cauliflower mosaic virus 35S promoter. Environmental Biosafety Research 3, 91–97.

Urbanek MO, Galka-Marciniak P, Olejniczak M, Krzyzosiak WJ. 2014. RNA imaging in living cells – methods and applications. RNA Biology 11, 1083–1095.

Wang H, Nakamura M, Zhao D, Nguyen CM, Yu C, Lo A, Daley T, Russa ML, Liu Y, Qi LS. 2019. Temporal-Spatial Visualization of Endogenous Chromosome Rearrangements in Living Cells. 734483.

Wang S, Su JH, Zhang F, Zhuang X. 2016. An RNA-aptamer-based two-color CRISPR labeling system. Scientific Reports 6, 26857.

Weijers D, Franke-van Dijk M, Vencken RJ, Quint A, Hooykaas P, Offringa R. 2001. An *Arabidopsis* Minute-like phenotype caused by a semi-dominant mutation in a RIBOSOMAL PROTEIN S5 gene. Development 128, 4289–4299.

Wesolowska N, Amariei FL, Rong YS. 2013. Clustering and protein dynamics of *Drosophila melanogaster* telomeres. Genetics 195, 381–391.

Winter D, Vinegar B, Nahal H, Ammar R, Wilson GV, Provart NJ. 2007. An “Electronic Fluorescent Pictograph” browser for exploring and analyzing large-scale biological data sets. PLoS ONE 2, e718.

Wu X, Mao S, Ying Y, Krueger CJ, Chen AK. 2019. Progress and Challenges for Live-cell Imaging of Genomic Loci Using CRISPR-based Platforms. Genomics, Proteomics & Bioinformatics 17, 119–128.

Xiang G, Zhang X, An C, Cheng C, Wang H. 2017. Temperature effect on CRISPR-Cas9 mediated genome editing. Journal of Genetics and Genomics 44, 199–205.

Xu W, Li K, Li S, Hou Q, Zhang Y, Liu K, Sun Q. 2020. The R-loop atlas of *Arabidopsis* development and responses to environmental stimuli. The Plant Cell 32, 888–903.

Zhang S, Song Z. 2017. Aio-Casilio: a robust CRISPR-Cas9-Pumilio system for chromosome labeling. Journal of Molecular Histology 48, 293–299.

Zuo J, Niu Q-W, Chua N-H. 2000. An estrogen receptor-based transactivator XVE mediates highly inducible gene expression in transgenic plants. The Plant Journal 24, 265–273.

